# WorldFlora: An R package for exact and fuzzy matching of plant names against the World Flora Online Taxonomic Backbone data

**DOI:** 10.1101/2020.02.02.930719

**Authors:** Roeland Kindt

## Abstract

**Premise of the study:** Standardization of plant names is a critical step in various fields of biology including biodiversity, biogeography and vegetation research. *WorldFlora* matches lists of plant names with a static copy from World Flora Online (WFO), an ongoing global effort of completing an online flora of all known vascular plants and bryophytes by 2020.

**Methods and results:** Based on direct and fuzzy matching, *WorldFlora* inserts matching cases from the WFO to a submitted data set of with taxa. Results of selecting the expected best single matches are presented for four data sets, including a working list of commercial timber tree species, a subset from *GlobalTreeSearch* and 2 data sets used in previous comparisons of software tools for correcting taxon names. The success rate of credible matches varied from 94.7 (568 taxa) to 99.9 (1740 taxa) percent.

**Conclusions:** *WorldFlora* offers a straightforward pipeline for semi-automatic plant name checking.

## INTRODUCTION

Scientific names of organisms and the higher groups in which they are classified are key identifiers of the world’s biodiversity (Rees, 2014). Assigning a species identify to an organism is essential in a wide array of disciplines including ecology, conservation and forestry (Tyrell, 2019). Removing synonyms from plant species lists is needed to predict the total number of vascular, seed and flowering plant species (Lughadha et al., 2016). Taxonomic uncertainty is one of the major gaps in plant occurrence data needed in global plant ecological, biogeographical and conservation applications (Meyer et al., 2016). Misspellings can lead to failures to retrieve data from global databases that encompass millions of species (Boyle et al., 2013). Matching alternative species names and resolving synonyms was for example essential in the development of the Agroforestry Species Switchboard, a portal that currently documents the presence of 172,395 plant species across 35 web-based information sources through 307,404 hyperlinks (Kindt et al., 2019). If not corrected, lack of standardization of names can lead to inflated estimates of species richness. For example, a revision of a checklist of 11,675 Amazonian tree species through a taxonomic vetting process resulted in an updated list of 10,071 species, representing a ‘loss’ of around 15% of taxa (ter Steege et al., 2019). Issues of misspellings and unresolved synonyms increases the risks of erroneous scientific conclusions, such as misidentification of medicinal applications in plants, with potentially serious consequences (Sharma et al., 2019).

Target 1 of the Global Strategy for Plant Conservation aims to complete an online flora of all known plants by 2020. To achieve this goal, the World Flora Council was formed, including 36 participating institutions (Miller and Ulate, 2017). Building on The Plant List (http://www.theplantlist.org) that became static in 2013, the World Flora Online’s (WFO) taxonomic backbone is actively curated by taxonomic specialists and Taxonomic Expert Networks (Palese et al., 2019). Static copies of the Taxonomic Backbone data are available from the WFO website (http://www.worldfloraonline.org/downloadData). The WorldFlora package matches a list of taxa with the WFO taxonomic backbone and was the only package known to do this at article submission.

## METHODS AND RESULTS

*WorldFlora* is an R package that matches a list of plant names with a static version of the Taxonomic Backbone data that can be downloaded from the World Flora Online portal (http://www.worldfloraonline.org/downloadData; examples presented herein used version 2019.05 of 17-May-2019, the most recent version available at article submission). Data sets can be imported in R in a wide variety of formats, with various available through the Graphical User Interface of the R-Commander (Fox, 2005). Exact matching with the WorldFlora::WFO.match function can be undertaken for the entire plant name or simultaneously for genus, species and infraspecifc levels. Fuzzy matching is implemented by WFO.match for the full plant name, calculating the Levenshtein Distance (LD) via R functions of base::agrep (a user can modify its argument of max.distance from the default 0.1) and utils::adist. The LD measures single character substitutions, insertions or deletions, each contributing a value of 1 to the total distance. The Taxamatch algorithm (Rees, 2014), also implemented in the Taxonomic Name Resolution Service (TNRS; Boyle et al., 2013), uses the Modified Damerau-Levenshtein distance where transpositions are given lower weights (2 in the example of ‘vecusilosus’ and ‘vesiculosus’) than the weights of substituting each character individually (4 in the example). Users can limit the number of matches to those with the smallest LD by setting argument Fuzzy.min = TRUE, the default setting. The WorldFlora::WFO.one function finds single matches for each submitted plant name by filtering records by accepted names and synonym status. Information about the rationale for selection is given in a separate column in the output. Where multiple candidates remained, the function selects the match with the smallest WFOID field. Successful matches by *WorldFlora* are limited to the scope of the World Flora Online of vascular plants and bryophytes, similar to software packages that use The Plant List such as Taxonstand (Cayuela *et al.*, 2012). Therefore, users ideally should not attempt to resolve names from organism groups that are not covered such as algae, fungi or lichens (Wagner, 2016).

Four data sets were used to check the performance of *WorldFlora* and to describe some of its features. The first is a random subset of 1,000 species selected from the GlobalTreeSearch (GTS) database (Beech et al., 2017; version 1.3 accessed from https://tools.bgci.org/global_tree_search.php). Of these, 957 species were matched directly (Table 1). The remaining 43 were matched by the fuzzy algorithm. Where several matches were retrieved, the first option of finding the single best match was by selecting the record with the smallest LD between the submitted and matched Authority (selecting the best author match is the default option for WorldFlora::WFO.one if the Authority variable is declared). This option resulted in 24 single matches, including 21 matches with a LD between authorities of zero (for example, the selection of *Bauhinia grandifolia* (Bong.) D.Dietr. and rejection of *Bauhinia grandifolia* Steud.; see Appendix S1). The remaining five single matches were based on not selecting a synonym name, with 4 of those having a LD between authorities of zero as by selecting *Xylosma intermedia* (Seem.) Griseb. and rejecting synonym *Xylosma intermedia* (Seem.) Triana & Planch. For 957 species, the name retrieved by *WorldFlora* was exactly the same as the submitted name (classified as ‘correct’ in Table 1). Among the names that were not matched exactly, three were spelling variants (*Callianthe sylvatica* vs. *Calanthe sylvatica, Euodia cuspidata* vs. *Evodia cuspidata* and *Guatteria moralesii* vs. *Guatteria moralesi*). Five names would have been matched if a scientist familiar with revision of the *Acacia* genus (Brummitt, 2004) would have substituted the genera *Senegalia* or *Vachellia* by *Acacia*, resulting in a tally of 965 (957 + 3 + 5; Table 1) ‘credible’ matches that can be achieved by *WorldFlora* in a typical supervised workflow. Of the 35 species without credible matches, 28 were partially matched at the genus level through the default option of argument ‘Fuzzy.one’ that searched for matches only for the first word, in case that the full name yielded no matches (Table2 provides an overview of various function arguments).

**Table 1.** Results of matching plant names from four data sets. The difference between correct and credible matches is clarified in the text. Details about access, data manipulation, outputs and R scripts are available from the appendices.

| Data set | GTS | CTTS | Wagner | SALVIAS |
| --- | --- | --- | --- | --- |
| Taxa | 1,000 | 1741 | 600 | 1,000 |
| Unique direct matches | 930 | 1513 | 0 | 694 |
| Several direct matches | 27 | 125 | 0 | 40 |
| Unique fuzzy matches | 41 | 100 | 491 | 232 |
| Several fuzzy matches | 2 | 3 | 41 | 22 |
| No match | 0 | 0 | 68 | 12 |
| Correct matches | 957 | 1728 | 500 | 951 |
| Credible matches | 965 | 1740 | 568 | 975 |
| Appendix | Appendix S1 | Appendix S2 | Appendix S3 | Appendix S4 |

Note: GTS = Global Tree Search data set; CCTS = Commercial Timber Tree Species data set; Wagner = testing data set from Wagner (2016); SALVIAS = testing data set from Boyle et al. (2013).

**Table 2.** Overview of some of the arguments of WorldFlora::WFO.match.

| Argument | Details |
| --- | --- |
| Authorship | If this variable is found in the submitted data, the result will include a column of 'Auth.dist' with the LD between the submitted and matched naming author. |
| acceptedNameUsageID.match | In the default setting, where the WFO includes a acceptedNameUsageID (typically indicating the accepted name for a synonym), then the matched details in the results will show the accepted name and the columns of 'Old.status', 'Old.ID' and 'Old.name' those of the first match. |
| Fuzzy.max | The maximum number of names matched by the fuzzy algorithm. |
| Fuzzy.min | In the default setting (TRUE), only the matches with the smallest LD are retained. |
| Fuzzy.two | 'Fuzzy.two' flags (TRUE/FALSE) whether the match was obtained for only the first 2 words of the submitted name |
| Fuzzy.one | 'Fuzzy.one' flags (TRUE/FALSE) whether the match was obtained for only the first word of the submitted name |
| spec.name.nonumber | 'Number.detected' flags (TRUE/FALSE) whether the submitted name contained a number. If that was the case, a match was searched only for the first word of the submitted name. |
| spec.name.tolower | Converts all characters of the submitted name to lower case, except the first character. |
| spec.name.nobrackets | 'Brackets.detected' flags (TRUE/FALSE) whether the submitted name contained a bracket. If that was the case, the function searched only for the part of the submitted name before the bracket. |

The second data set used as a case study was a working list of 1,741 commercial timber tree species (CTTS; Mark et al., 2014). Names were matched directly for 1,638 taxa and by fuzzy matching for the remaining 103 names (Table 1). Of the fuzzy matches, 88 had a LD of 1 due to a space character at the end of the submitted name. The WFO.one function used the criterion of the smallest WFOID for 14 names (for example selecting wfo-0000913932: *Scottellia coriacea* A.Chev. ex Hutch. & Dalziel and rejecting wfo-0000913938: *Scottellia coriacea* A.Chev.), but in only one case were the candidate species different (selecting synonym wfo-0000416717: *Ulmus glabra* and rejecting synonym wfo-0000475766: *Planera aquatica* for the submitted *Ulmus campestris*). It was confirmed for 28 species that where the original data set had included a synonym name between brackets, that this was indeed the accepted synonym name (for example, *Dysoxylum euphlebium* to be a synonym of *Dysoxylum alliaceum*). Among the 13 species where the submitted and retrieved name was not exactly the same (not ‘correct’ as in Table 1), 12 were spelling variants such as *Pinus englemannii* vs. *Pinus engelmannii* or *Sequoiadendron gigantum* vs. *Sequoiadendron giganteum. Populus canescens* was matched by its hybrid name *Populus ×canescens*, correctly reflecting that the species is now accepted to be a hybrid of *Populus alba × Populus tremula* (Tutin et al., 1993). The only species without a credible match was *Upuna borneensis* (Randi et al., 2019) that was matched by *Upuna boreensis*.

The third data set that was tested was a combination of three sub data sets initially created to compare different software packages for correcting plant names (‘Wagner’, Wagner, 2016). The argument Fuzzy.max was increased to 2,500 because an initial run of WFO.match could not resolve many of the names submitted at genus rank. As the testing procedure involved deleting the last character from a list of species names, all the matches were fuzzy, as might be expected (Table 1). One hundred of the names were not identical to the expected names (not ‘correct’ in Table 1). Of these, 60 were names of algae, fungi and lichen species outside the scope of World Flora Online and its predecessor The Plant List. Five of these names were spelling variants, one was an interspecies hybrid (*Carex acuta* × *elata*) and two were matched as varieties rather than the submitted subspecies names (*Keckiella antirrhinoides* var. *microphylla* for *Keckiella antirrhinoides* subsp. *microphylla, Saxifraga adscendens* var. *oregonensis* for *Saxifraga adscendens* var. *oregonensis*). Reasons that no acceptable matches were found for the remaining 32 species included 7 names where the submitted family is not included in the WFO (Aceraceae, Najadaceae, Punicaceae, Taccaceae, Theophrastaceae, Tiliacae and Vittariaceae). None of these families were retained in the fourth update of the Angiosperm Phylogeny Group (2016) classification of orders and families of angiosperms. Seven were names where the number of fuzzy matches was too high to retrieve the genus name, while there were nine names where a submitted hybrid was matched by a non-hybrid name. In three cases, mismatches resulted from selecting the record with the smallest ID from otherwise valid fuzzy alternatives (selecting *Lomatia* R.Br. [Proteaceae] and not *Lomatium* Raf. [Apiaceae], *Placea* instead of Poaceae and *Sabicea* instead of Sabiaceae). With the overall matching success of 87.6 percent (219 out of 250 species) for spellchecking tests across different taxonomic ranks, WorldFlora performed better than the 74 percent achieved by the Global Names Resolver (GNR), which was the best software in the comparisons undertaken by Wagner (2016). With 99.6 percent correct matches in comparisons across different geographic regions (only *Leptinella conjuncta* was not matched from the 250 submitted names), WorldFlora also marginally outperformed GNR as the best software, with GNR giving 99.2 percent correct matches in the earlier comparison (Wagner, 2016).

The fourth data set that was analysed, ‘SALVIAS’, is a list of 1,000 plant names that was used earlier in a comparison of TNRS with other online tools for automated standardization of plant names (Boyle et al., 2013). The same data set was used in a more recent evaluation study (Sharma et al., 2019). This data set offers various challenges to plant name checking, such as inclusion of names in capital letters. This particular challenge is handled by WFO.match by option spec.name.tolower=TRUE, whereby submitted names are converted to lower case, except for the first character. The challenge of handling semi-standardized qualifiers that are used in plant names, such as ‘cf.’ that indicates that not all of the diagnostic characters correspond to a given species, or ‘aff.’ that indicates that a specimen has some affinity but is not identical to a known species, is handled by deleting these characters from the name. Taxamatch (Rees, 2014) and TNRS (Boyle et al., 2013) use a similar approach. The default list of qualifiers used by WFO.match was derived from descriptions provided by Sigovini et al. (2016), also including qualifiers such as ‘sp.’, ‘indet.’ and ‘nom. nov.’. As WFO.match expects that the submitted plant name does not include the naming authority, when no matches are found with the submitted name that included the authority, a search is done for the first two words of the submitted name now expected to correspond to the genus and species names (default option for argument Fuzzy.two). If the search still does not find a name match, then a match is attempted for the first word only (default option for argument Fuzzy.one). Names that contain brackets, possibly at the beginning of the authority name, are stripped from the entire part starting with the bracket (default option of spec.name.nobrackets). Names that contain numbers are searched only for the first word, with the remaining part of the submitted name suspected to correspond to unidentified species (default option of spec.name.nonumber).

An initial run of WorldFlora.match with SALVIAS yielded 25 names with no match. Visual inspection of these unmatched names revealed cases where a qualifier ‘cf’ was used instead of the standard ‘cf.’ or ‘aff’ instead of ‘aff.’. Simulating how an actual semi-automatic pipeline of plant name checking would work, incomplete qualifiers were replaced by the correct qualifier that is recognized in the argument of ‘sub.pattern’. Likewise, argument Fuzzy.max was increased to 2,000, as the output of WorldFlora.match indicated a series of names where the number of fuzzy matches was above the default 250, with a maximum of 1,974 fuzzy matches for the submitted ‘Miconia’. The second run of WorldFlora.match with the modified names and arguments resulted in 734 directly matched names and 254 names with fuzzy matches (Table 1). Directly comparing the final subset of names obtained via WorldFlora.one with those obtained by TNRS (http://tnrs.iplantcollaborative.org/TNRSapp.html; accessed on 12-12-2019 with default settings) showed 951 identical names (‘correct’ in Table 1). Among the 49 names that were not identical, eight were spelling variants such as *Commiphora laxecymigera* vs. *Commiphora laxicymigera*. For 16 of the names that were not identical, WorldFlora resulted in a more credible match than TNRS. For example, the submitted Asteraceae and Fabaceae families were correctly matched by WorldFlora, but matched incorrectly by the Asteliaceae or Fagaceae by the TNRS. For example, TNRS provided a partial match of the submitted *Solanum schlechtendali* to *Solanum*, whereas WorldFlora correctly matched *Solanum schlechtendalianum*. Accepting these credible matches by WorldFlora resulted in the correct matching of 975 names, a number close to the 980 correct matches reported for TNRS (Boyle et al., 2013). The number of matches is above the 950 of Solr-Plant for the same data set, which performed second best among TNRS, Solr-Plant, Plantminer and GNResolver (Sharma et al., 2019).

## CONCLUSIONS

Analysis of the four data sets showed that the new package WorldFlora matched plants names correctly for the majority of submitted names, with success rates comparable to and sometimes better than alternative software packages such as the Taxonomic Name Resolution Service or Solr-Plant. Users are advised not to blindly accept results, however, as limitations in plant name checking software and spelling mistakes in reference databases can result in incorrect matches (Wagner, 2016). Users are therefore especially advised to screen the output of the WorldFlora::WFO.match and WorldFlora::WFO.one functions for information that could indicate possible mismatches, such as a large LD between the submitted and matched plant names, or the justification of a single match based on the smallest WFOID. Where possible, users are further advised to compare results obtained between different applications, for example, by submitting names both to WorldFlora and TNRS, first checking where both software packages agree on the accepted plant name. As World Flora Online provides a further development of The Plant List that became static in 2013, matching of plant names with *WorldFlora* is expected to provide results that are more up-to-date than software packages that rely on The Plant List, such as Taxonstand (Cayuela et al., 2012). To benefit maximally from the current state of knowledge on plant names, users should ensure that they use the most recent version of the World Flora Online backbone database from the website. Since naming authorities are not part of the plant name that is checked, users ideally should submit the plant name and the authority in separate fields to reduce mismatches.

## ACKNOWLEDGMENTS

RK thanks Ian Dawson (ICRAF) for reviewing the article prior to submission, and XX anonymous reviewers for further improving this manuscript. RK thanks the CGIAR Research Programs on Forests, Trees and Agroforestry (supported by the CGIAR Trust Fund) and the Provision of Adequate Tree Seed Portfolios project (supported by the Royal Norwegian Embassy in Ethiopia and NICFI) for supporting the author’s time to contribute.

## DATA AVAILABILITY

The *WorldFlora* package is published under the GNU General Public License (Version 2). The software and related documentation are available for free download from the Comprehensive R Archive Network (https://cran.r-project.org/package=WorldFlora).

## APPENDIX

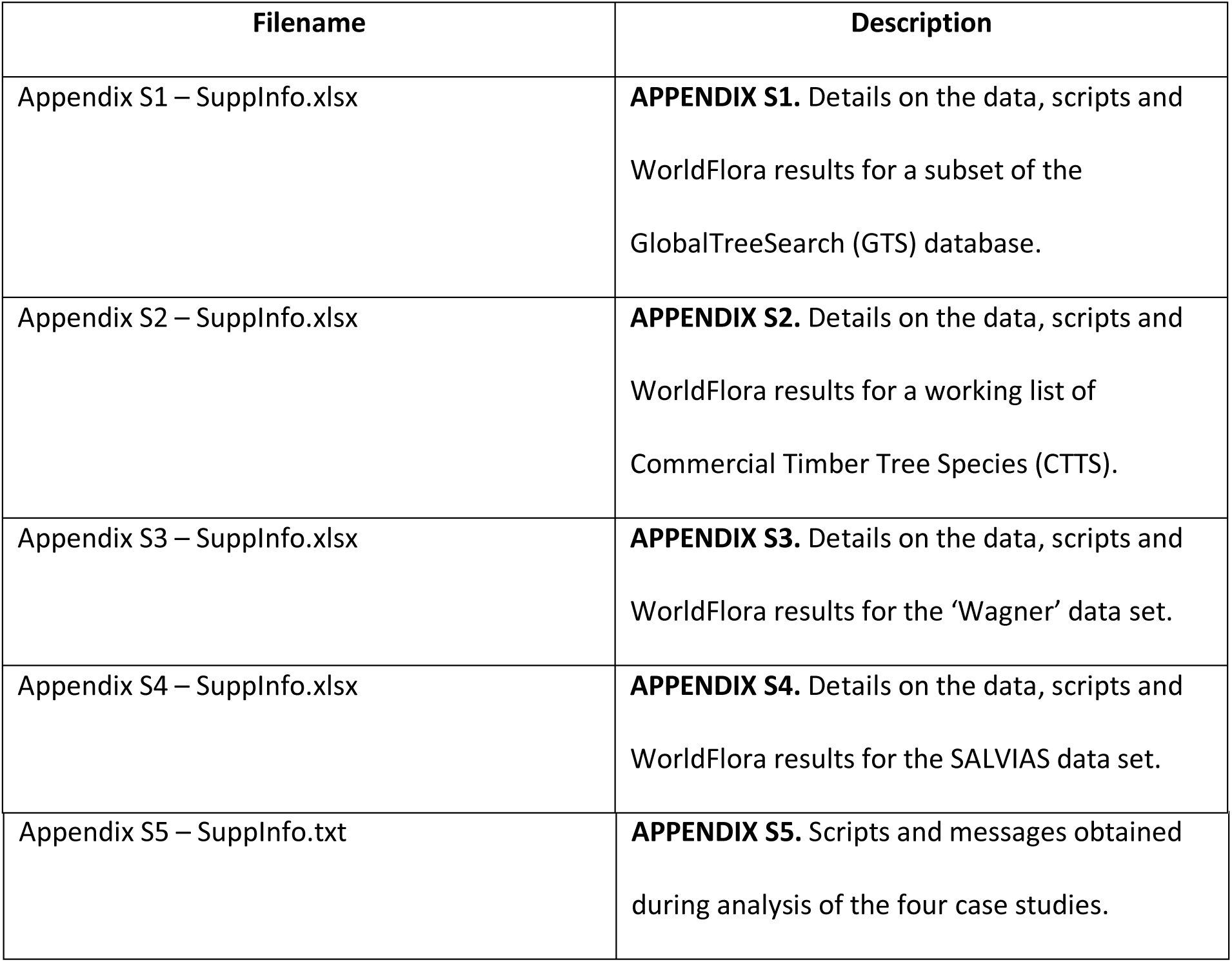

## LITERATURE CITED

Beech, E., M. Rivers, S. Oldfield, and P. P. Smith. 2017. GlobalTreeSearch: the first complete global database of tree species and country distributions. Journal of Sustainable Forestry 36: 454–489.

Boyle, B., N. Hopkins, Z. Lu, J. A. R. Garay, D. Mozzherin, T. Rees, N. Matasci et al. 2013. The taxonomic name resolution service: an online tool for automated standardization of plant names. BMC Bioinformatics 14: 16.

Brummitt, R. K. 2004. Report of the Committee for Spermatophyta: 55. Proposal 1584 on *Acacia*. TAXON 53: 826–829.

Cayuela, L., I. Granzow-de la Cerda, F. S. Albuquerque, and D. J. Golicher. 2012. Taxonstand: An r package for species names standardisation in vegetation databases. Methods in Ecology and Evolution 3: 1078–1083

Fox, J. 2005. The R Commander: A Basic Statistics Graphical User Interface to R. Journal of Statistical Software 14: 1–42.

Hackett, R. A., M. W. Belitz, E. E. Gilbert, and A. K. Monfils. 2019. A database management workflow of biodiversity data from the field to data users. Applications in Plant Sciences 7: e11310.

Kindt, R., I. John, J. Ordonez, I. Dawson, J.-P. B. Lillesø, A. Muchugi, L. Graudal, and R. Jamnadass. 2019. Agroforestry Species Switchboard: a synthesis of information sources to support tree research and development activities. Version 2.0. Website http://www.worldagroforestry.org/products/switchboard [accessed 1 February 2020]

Lughadha, E. N., R. Govaerts, I. Belyaeva, N. Black, H. Lindon, R. Allkin, R. E. Magill, and N. Nicolson. 2016. Counting counts: revised estimates of numbers of accepted species of flowering plants, seed plants, vascular plants and land plants with a review of other recent estimates. Phytotaxa 272: 82–88.

Mark, J., A. C. Newton, S. Oldfield, and M. Rivers. 2014. The International Timber Trade: A working list of commercial timber tree species. Botanic Gardens Conservation International, Richmond, United Kingdom.

Miller, C. and W. Ulate. 2017. World Flora Online Project: An online flora of all known plants. Proceedings of TDWG 1: e20529.

Palese, R., C. Boillat, and P.-A. Loizeau. 2019. World Flora Online (WFO) – Quality control workflow for an evolving taxonomic backbone. Biodiversity Information Science and Standards 3: e35307.

Randi, A., V. Bodos, and J. Pereira, J. 2019. Upuna borneensis. The IUCN Red List of Threatened Species 2019: e.T33148A68075816.

Rees, T. 2014. Taxamatch, an Algorithm for Near (‘Fuzzy’) Matching of Scientific Names in Taxonomic Databases. PLOS ONE 9: e107510.

Sharma, V., M. I. Restrepo, and I. N. Sarkar. 2019. Solr-Plant: efficient extraction of plant names from text. BMC Bioinformatics 20: 263

Sigovini, M., E. Keppel, and D. Tagliapietra. 2016. Open Nomenclature in the biodiversity era. Methods in Ecology and Evolution 7: 1217–1225.

ter Steege, H., S. M. de Oliveira, N. C. A. Pitman, D. Sabatier, A. Antonelli, J. E. G. Andino, G. A. Aymard, and R. F. Salomao. 2019. Scientific Reports 9: 3501.

The Angiosperm Phylogeny Group, M. W. Chase, M. J. M. Christenhusz, M. F. Fay, J. W. Byng, W. S. Judd, D. E. Soltis, et al. 2016. An update of the Angiosperm Phylogeny Group classification for the orders and families of flowering plants: APG IV, Botanical Journal of the Linnean Society 181: 1–20.,

Tutin, T.G., N. A. Burges, A. O. Chater, J. R. Edmondson, V. H. Heywood, D. M. Moore, D. H. Valentine, et al. 1993). Flora Europaea Volume 1: PSILOTACEAE TO PLATANACEAE. Cambridge University Press, Cambridge, United Kingdom.

Tyrell, C. D. 2019. A method to implement continuous characters in digital identification keys that estimates the probability of an annotation. Applications in Plant Sciences 7: e1247

Wagner, V. 2016. A review of software tools for spell-checking taxon names in vegetation databases. Journal of Vegetation Science 27: 1323–1327.

